# Differential protein metabolism and regeneration in hypertrophic diaphragm and atrophic gastrocnemius muscles in hibernating Daurian ground squirrels

**DOI:** 10.1101/793752

**Authors:** Xia Yan, Xuli Gao, Xin Peng, Jie Zhang, Xiufeng Ma, Yanhong Wei, Huiping Wang, Yunfang Gao, Hui Chang

## Abstract

Whether differences in regulation of protein metabolism and regeneration are involved in the different phenotypic adaptation mechanisms of muscle hypertrophy and atrophy in hibernators? Two fast-type muscles (diaphragm and gastrocnemius) in summer active and hibernating Daurian ground squirrels were selected to detect changes in cross-sectional area (CSA), fiber type distribution, and protein expression indicative of protein synthesis metabolism (protein expression of P-Akt, P-mTORC1, P-S6K1, and P-4E-BP1), protein degradation metabolism (MuRF1, atrogin-1, calpain-1, calpain-2, calpastatin, desmin, troponin T, Beclin1, and LC3-II), and muscle regeneration (MyoD, myogenin, and myostatin). Results showed the CSA of the diaphragm muscle increased significantly by 26.1%, whereas the CSA of the gastrocnemius muscle decreased significantly by 20.4% in the hibernation group compared with the summer active group. Both muscles displayed a significant fast-to-slow fiber-type transition in hibernation. Our study further indicated that increased protein synthesis, decreased protein degradation, and increased muscle regeneration potential contributed to diaphragm muscle hypertrophy, whereas decreased protein synthesis, increased protein degradation, and decreased muscle regeneration potential contributed to gastrocnemius muscle atrophy. In conclusion, the differences in muscle regeneration and regulatory pattern of protein metabolism may contribute to the different adaptive changes observed in the diaphragm and gastrocnemius muscles of ground squirrels.

## Introduction

Although hibernating mammals are recognized as models of disuse-resistant skeletal muscle atrophy (Bodine 2013, Ivakine and Cohn 2014, Lee et al. 2008, Reilly and Franklin 2016), not all types of skeletal muscle maintain constant quality during long periods of hibernation. Our and other previous studies have found that different types of skeletal muscles exhibit different degrees of atrophy during mammalian hibernation (Cotton 2016, Zhang et al. 2019). The diaphragm is a fast-type respiratory muscle. Inactivity caused by mechanical ventilation can lead to diaphragm atrophy (Powers et al. 2013). However, although breathing in hibernating mammals is considerably inhibited during hibernation (Larson et al. 2014, Staples and Brown 2008), such a condition does not induce diaphragm atrophy, but rather hypertrophy. For example, previous studies have reported increases in diaphragm muscle weight of 31.1% and 19% in hibernating Syrian hamsters (*Mesocricetus auratus*) (Deveci and Egginton 2002) and golden-mantled ground squirrels (*Callospermophilus lateralis*), respectively (Reid et al. 1995, Rourke et al. 2004). However, in the fast-type gastrocnemius muscle, hindlimb unloading in non-hibernating rats resulted in a 42% decrease in muscle weight and 34%–46% decrease in the cross-sectional area (CSA) of slow and fast muscle fibers (Hu et al. 2017). In addition, muscle weight in the gastrocnemius muscle also showed a decrease in small hibernating rodents (8.1%–29.8%), but to a lesser degree and with less atrophy than that found in non-hibernators (Cotton 2016).

Skeletal muscles, as highly plastic tissue, undergo certain adaptive changes in structure and function when the environment changes (Blaauw et al. 2013). Muscle size is controlled by the balance between protein synthesis and degradation. Muscle hypertrophy occurs when synthesis is greater than degradation; in contrast, muscle atrophy occurs when degradation is greater than synthesis (Romanello and Sandri 2016). The Akt-mTOR signaling pathway plays a key role in the initiation of protein synthesis in skeletal muscle under the physiological state (Bodine et al. 2001). Activated mTORC1 (mammalian target of rapamycin in complex 1) phosphorylates downstream targets S6K1 (70 kDa ribosomal protein S6 kinase 1) and 4E-BP1 (eukaryotic initiation factor 4E-binding protein 1), resulting in increased protein synthesis (Gordon et al. 2013, Liu et al. 2013). During hibernation, however, the phosphorylation and activity of Akt and mTOR can be modified. For example, both are decreased in the thigh and femoral muscles of thirteen-lined ground squirrels (*Ictidomys tridecemlineatus*) during hibernation and in the pectoral muscles of greater tube-nosed bats (*Murina leucogaster ognevi*) during torpor (Cai et al. 2004, Lee et al. 2010, Wu and Storey 2012). Furthermore, the expression of phosphorylated 4E-BP is also decreased in the thigh muscles of thirteen-lined ground squirrels during hibernation (Wu and Storey 2012).

Protein degradation of skeletal muscles mainly includes the calpain degradation, ubiquitin-proteasome, and autophagy-lysosome pathways (Scicchitano et al. 2015). Calpains are the promoters of muscle protein degradation, which can release myofibrillar proteins from sarcomeres. The released myofibrillar proteins can be further ubiquitized and transported to proteasomes for degradation (Bartoli and Richard 2005, Huang and Forsberg 1998). Muscle RING finger 1 (MuRF1) and muscle atrophy F-box (MAFbx)/atrogin-1 are ubiquitin protein ligases and important molecular markers for evaluating skeletal muscle atrophy (Foletta et al. 2011). The ubiquitin proteasome system depends on multiple proteasomes, of which 26S proteasome can cleave the target protein as peptides (Chen et al. 2015). 26S proteasome is composed of 20S proteasome which mediates the activity of chymotrypsin and trypsin, and plays an important role in ubiquitin protease system (Souza et al. 2014). We previously found that calpain-1 and calpain-2 protein expression is decreased in the soleus muscle, but unchanged in the extensor digitorum longus (EDL) muscle in Daurian ground squirrels during hibernation, whereas the expression of calpastatin, an endogenous inhibitor of calpains, is increased in both (Chang et al. 2018b). Calpain can degrade substrates such as desmin, troponin T, troponin I, titin, alpha-fodrin and alpha-actinin, (Barta et al. 2005). Desmin and troponin T are more sensitive to calpain than other substrates (Li et al. 2017). Desmin is located around the Z-discs and connects the adjacent Z-discs to the sarcolemma and the nucleus (Capetanaki et al. 1997). Troponin T plays an important role in regulating protein system, which can bind to tropomyosin to form a complex (Perry 1998). Furthermore, various stress conditions (including fasting, oxidative stress, and hypoxia) can induce autophagy as an adaptive physiological response to restore cell metabolism. Increased autophagy is a survival-promoting mechanism rather than a death-promoting mechanism (He et al. 2012, Levine and Yuan 2005, Moresi et al. 2012), and is an important pro-survival mechanism in myocardial hibernation (May et al. 2008). However, previous study has indicated that autophagy is inhibited in fast muscles, including quadriceps and anterior tibial muscles, of hibernating thirteen-lined ground squirrels (Andres-Mateos et al. 2013). Therefore, we hypothesize that the opposite adaptabilities of skeletal muscles in hibernating Daurian ground squirrels, that is, hypertrophy and atrophy, are due to changes in the balance between protein synthesis and degradation.

Muscle regeneration includes an increase in protein synthesis (hypertrophy) and proliferation of muscle satellite cells (hyperplasia) and is a repair mechanism of muscle injury (Zanou and Gailly 2013). Previous studies have shown that the regeneration potential of disused skeletal muscle is changed (Suetta et al. 2013, Wall et al. 2014). MyoD (myogenic differentiation) and myogenin are myogenic regulatory factors (MRFs). MyoD is a molecular marker of muscle satellite cells activation into muscle progenitor cells (Zanou and Gailly 2013), and myogenin plays a key role in the fusion of myoblasts to form myotubes (Hasty et al. 1993). Even though the number of satellite cells decreases during mechanical unloading, the expression of markers of satellite cell activation increases (Arentson-Lantz et al. 2016, Guitart et al. 2018). Myostatin, a transforming growth factor-beta (TGFβ) family member, is a negative regulator of the proliferation and differentiation of muscle satellite cells. Inactivation of myostatin can lead to skeletal muscle hypertrophy, whereas overexpression can lead to muscle atrophy (Langley et al. 2002, McFarlane et al. 2011, Rodriguez et al. 2014). Myostatin not only regulates the growth and size of skeletal muscle, but also interacts with the Akt-mTOR pathway to regulate protein synthesis (Schiaffino et al. 2013, Trendelenburg et al. 2009). So far, however, contradictions still exist regarding whether skeletal muscle regeneration is involved in maintaining muscle quality in hibernating small mammals. For example, previous studies have reported no atrophy in muscles ablated of satellite cells in hibernating thirteen-lined ground squirrels (Andres-Mateos et al. 2012), but also reported that muscle satellite cells are not quiescent and actually increase during hibernation (Brooks et al. 2015). Therefore, in the current study, we attempted to clarify any differences in regeneration between two types of skeletal muscle with different adaptive changes, and whether regeneration plays a differential regulatory role in muscle maintenance in hibernating Daurian ground squirrels.

In this study, we examined two fast-type muscles (diaphragm and gastrocnemius) that exhibit different adaptative changes during hibernation (hypertrophic and atrophic adaptation, respectively) in Daurian ground squirrels. To elucidate the differential regulation of protein metabolism and muscle regeneration in the two types of skeletal muscle, we measured the CSA, fiber type distribution, protein synthesis metabolism (protein expression of P-Akt, P-mTORC1, P-S6K1, and P-4E-BP1), protein degradation metabolism (MuRF1, atrogin-1, calpain-1, calpain-2, calpastatin, desmin, troponin T, Beclin1 and LC3-II), proteasome activity, and muscle regeneration (MyoD, myogenin and myostatin) in the diaphragm and gastrocnemius muscles of summer active (SA) and hibernating (HIB) Daurian ground squirrels.

## Methods

### Acquisition and use of animals

The animals’ acquisition and use were approved by the Northwest University Ethics Committee. As described by our laboratory previously (Chang et al. 2018a, Chang et al. 2018b), sixteen Daurian ground squirrels of both sexes were captured from the Weinan plain in the Shaanxi province of China and fed with water and rat chow *ad libitum*. The ground squirrels were moved to a cold room at 4-6°C when they entered torpor in November with 1 individual per cage. The ground squirrels used in this experiment were in a natural hibernation state. The time of entering torpor was judged by putting sawdust on the back of each individual and reducing a body temperature (T_b_) below 9°C. T_b_ was measured by thermal imaging using a visual thermometer (Fluke VT04 Visual IR Thermometer, USA). According to our observations, the ground squirrels did not eat and drink after entering torpor state. Moreover, most ground squirrels recovered to hibernation state after 1-2 days of interbout arousals. The ground squirrels were divided into 2 groups randomly (*n* = 8, with 5 females and 3 males in each group): (1) summer active (SA): active ground squirrels in July with a T_b_ of 36-38°C; (2) hibernation (HIB): ground squirrels experiencing two months of hibernation with T_b_ maintained at 5-8°C and were sampled in torpor state.

### Muscle collection

After the body weight was recorded, the ground squirrels were anesthetized with 90 mg/kg sodium pentobarbital intraperitoneally. Then the diaphragm and gastrocnemius muscles from both legs were separated and removed. Subsequently, the muscle samples were put into liquid nitrogen and stored for the follow-up experiments. The animals were sacrificed by an overdose injection of sodium pentobarbital while finishing surgical intervention.

### Immunofluorescent analysis

The muscle fiber cross-sectional area (CSA) and fiber type composition were measured by immunofluorescent analysis as described by our laboratory previously (Chang et al. 2018b). Briefly, 5 mm of the mid-belly of lateral gastrocnemius muscle was cut into 10-μm thick frozen muscle cross-sections at −20°C with a cryostat (Leica, Wetzlar, CM1850, Germany). Because of the uneven thickness of the diaphragm, the fixed geometric position of the muscle strips was cut for slicing (see the yellow box in **Figure 1A**). Then the slides with sections were fixed in 4% paraformaldehyde for 30 min, and subsequently permeabilized in 0.1% Triton X-100 (dissolve in PBS) for 30 min. The slides were then blocked with 1% bovine serum albumin (BSA) in PBS for 60 min at room temperature, and immediately incubated at 4°C overnight with the anti-laminin rabbit polyclonal antibody (1:200, Boster, BA1761-1) to visualize the myofiber interstitial tissue and the anti-skeletal slow myosin mouse monoclonal antibody (1:200, Boster, BM1533) to visualize the type I MHC in both muscles. The slides were rinsed twice in PBS and incubated with Alexa Fluor 647-labeled IgG secondary antibody (1:200; Thermo Fisher Scientific, #A21245) and FITC-labeled IgG secondary antibody (1:200; Sigma, F1010) for 60 min. The staining method of anti-MYH1 (IIx) mouse antibody (1:400, Abcam, ab127539) was the same as that of anti-skeletal slow myosin mouse monoclonal antibody, only the first antibodies were different, with anti-MYH1 (IIx) mouse antibody followed by FITC-labeled IgG secondary antibody. Anti-MYH2 (IIa) rabbit antibody and anti-laminin rabbit polyclonal antibody need to be stained separately. First, anti-MYH2 (IIa) rabbit antibody (1:200, Abcam, ab124937) was incubated overnight at 4°C and followed by Alexa Fluor 488-labeled anti-rabbit antibody (Thermo Fisher Scientific, A-21235). Next, anti-laminin rabbit polyclonal antibody was incubated overnight at 4°C and followed by Alexa Fluor 647-labeled anti-rabbit antibody. The staining method of anti-MYH4 (IIb) rabbit antibody (1:100, Santa Cruz Biotechnology, #34820) was the same as that of anti-MYH2 (IIa) rabbit antibody, only the first antibodies were different, with anti-MYH4 (IIb) rabbit antibody followed by Alexa Fluor 488-labeled anti-rabbit antibody. Photographs were acquired by using confocal microscopy (Olympus, Osaka, Japan) at an objective magnification of 40×. Image -Pro Plus 6.0 software was used to measure the CSA of at least 10 different visual fields of each sample.

**Figure 1.**
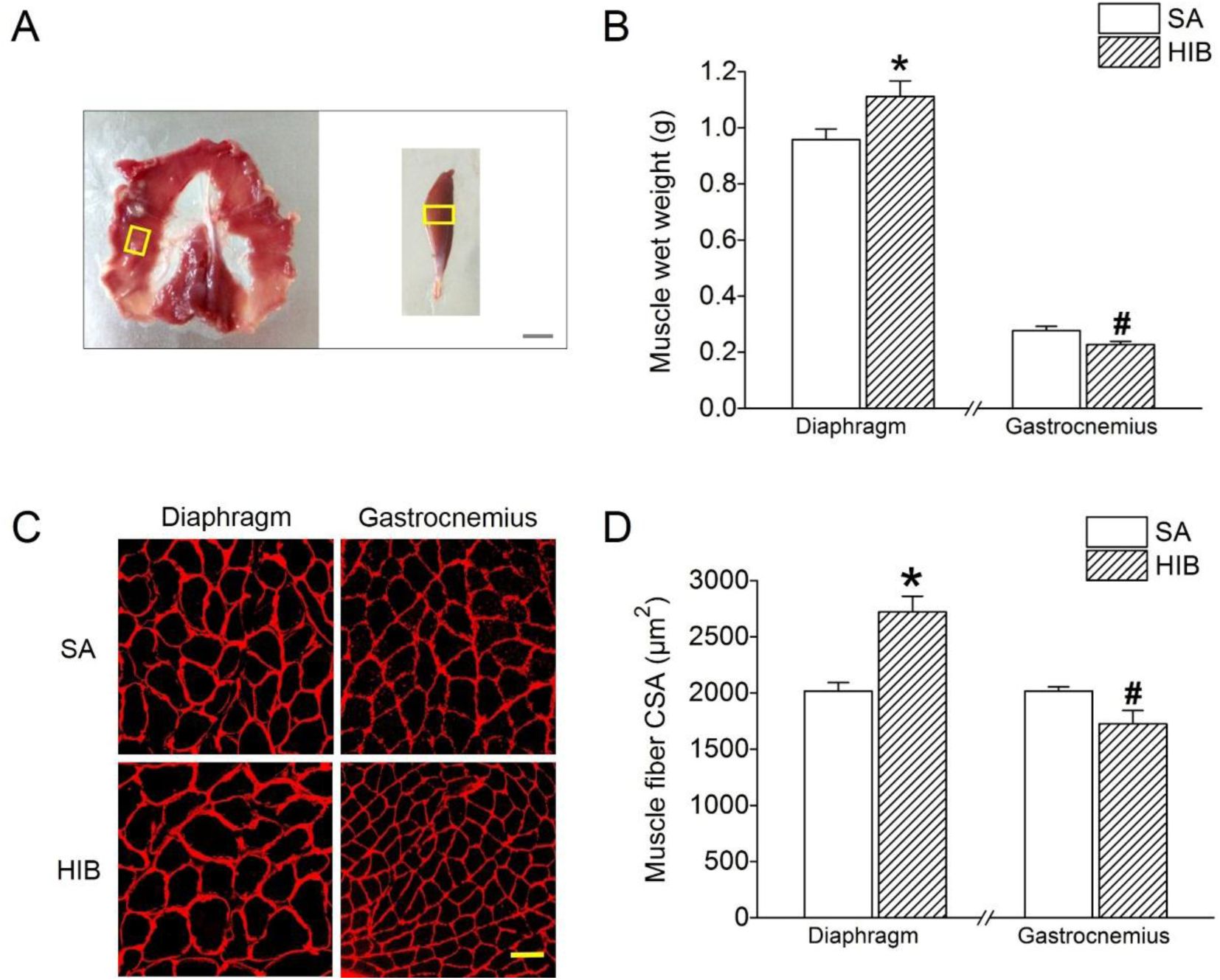
Muscle wet weight and fiber cross-sectional area (CSA) in diaphragm and gastrocnemius muscles. **(A)** Photos of diaphragm (left) and gastrocnemius (right) muscles. Scale bar = 4 mm. The yellow box shows the fixed geometric position for frozen section. **(B)** Muscle wet weight (g) in different groups. **(C)** Representative images showing cross-sections in diaphragm and gastrocnemius muscles. Red represents laminin stain of myofiber interstitial tissue. Scale bar = 50 μm. **(D)** Bar graph showing the changes of muscle fiber CSA. SA: summer active ground squirrels, HIB: hibernating ground squirrels. Data represents mean ± SEM, *n* = 8. \**P* < 0.05, compared with SA in diaphragm muscle. ^#^*P* < 0.05, compared with SA in gastrocnemius muscle.

### Proteasome activity

The chymotrypsin and trypsin activities of diaphragm and gastrocnemius muscles were determined by ultraviolet spectrophotometry according to the instructions of chymotrypsin assay kit (Nanjing Jiancheng Bioengineering Institute, A080-3-1) and trypsin assay kit (Nanjing Jiancheng Bioengineering Institute, A080-2-2).

### Western blots

As described previously by our lab (Chang et al. 2016, Chang et al. 2018b), the total protein was extracted from the frozen diaphragm and gastrocnemius muscles of ground squirrels by homogenization and put into a sample buffer (pH 6.8, 100 mM Tris, 4% SDS, 5% glycerol, 5% 2-β-mercaptoethanol, and bromophenol blue). Then the muscle protein extracts were resolved by SDS-PAGE using Laemmli gels (10%) with the 1% 2,2,2-Trichloroethanol (TCE) (aladdin, JI522028, China). After electrophoresis, the total proteins bands are visualized by putting the gel on the UV transilluminator and irradiating the gel for 2 min, and Syngene G:BOX system (Syngene, Frederick, MD) was used to take photographs of the gel. Then the protein gel was transferred electrically to PVDF membranes (0.45 μm) using a Bio-Rad semi-dry transfer apparatus. The blotted PVDF membranes were incubated with 1% BSA in TBS (Tris-buffered saline; 50 mM Tris– HCl, 150 mM NaCl, pH 7.5) and anti-skeletal slow myosin mouse monoclonal antibody (1:500, Boster, BM1533), anti-MYH2 (IIa) rabbit antibody (1:1000, Abcam, ab124937), anti-MYH4 (IIb) rabbit antibody (1:1000, Santa Cruz Biotechnology, #34820), anti-MYH1 (IIx) mouse antibody (1:1000, Abcam, ab127539), P-Akt (Ser473) (1:1,000, abcam, 81283), P-mTORC1 (Ser2448)(1:1,000, sigma, 4504476), P-S6K1 (Thr389) (1:1,000, Cell Signaling Technology (CST), 9205S), P-4E-BP1 (Thr37/46)(1:1,000, CST, 2855S), calpain-1 (1:1,000, CST, 2556S), calpain-2 (1:1,000, CST, 2539S), calpastatin (1:1,000, CST, 4146S), desmin (1:1,000, CST, 4024S), troponin T (1:1,000, Sigma, T6277), MuRF1 (1:1000, Abcam, ab172479), atrogin-1 (1:1,000, Proteintech, 12866-1-AP), Beclin1 (1:1000, CST, 3738), LC3 (1:1000, abcam, ab48394), MyoD (1:1,000, proteintech, 18943-1-AP), myogenin (1:1,000, abcam, 124800) and myostatin (1:1,000, proteintech, 19142-1-AP) antibodies, respectively, in TBS containing 0.1% BSA at 4°C overnight. Then the membrane was washed with TBST for 4 times (10 minutes/time) and incubated with HRP-conjugated anti-mouse secondary antibody (1:10000, Thermo Fisher Scientific, A28177) or HRP-conjugated anti-rabbit secondary antibody (1:5000, Thermo Fisher Scientific, A27036) for 2 h at room temperature. Then the PVDF membrane was washed for 3 times (10 min/time). The fluorescent band were visualized using enhanced chemiluminescence reagents (Thermo Fisher Scientific, NCI5079). The NIH Image J software was used to carry out quantification analysis. The density of immunoblot band in each individual lane was standardized by the summed densities from a group of total protein bands in the same lane. This method of standardizing by total protein has been proved to be more accurate and appropriate in comparison with standardizing by housekeeping proteins such as GAPDH, actin and tubulin (Vigelso et al. 2015, Zhang et al. 2016b).

### Statistical analyses

All data were analyzed using SPSS 19.0 and expressed as means ±SEM. An independent-samples *t*-test was used to determine the significant differences between SA and HIB ground squirrels. A value of *P* < 0.05 was considered to be statistically significant.

## Results

### Body weight and muscle wet weight

As shown in **Table 1**, the body weight of HIB group was significantly reduced by 11% (*P* < 0.05) as compared with SA group at experiment time.

**Table 1.** Body weight (BW) for all the groups.

|  | BW before hibernation (g) | BW at experiment time (g) |
| --- | --- | --- |
| SA | 288.3 ± 7.9 | 277.0 ± 16.2 |
| HIB | 277.0 ± 16.2 | 247.8 ± 21.6* |

SA: summer active ground squirrels, HIB: hibernating ground squirrels. Data represent mean ± SEM, $n = 8$ . \* $P < 0.05$ compared with SA group.

Compared with SA group, the muscle weight of the gastrocnemius muscle in the HIB group was significantly reduced by 18% (*P* < 0.05), while the muscle weight of the diaphragm muscle was significantly increased by 14% (*P* < 0.05) **(Figure 1B)**.

### Fiber CSA and fiber type distribution

The CSA and fiber type distribution of the diaphragm and gastrocnemius (lateral) muscle fibers were measured by immunofluorescent staining. Compared with the SA group, the CSA of the diaphragm muscle increased significantly (26.1%, *P* < 0.05), whereas that of the gastrocnemius muscle decreased significantly (20.4%, *P* < 0.05) in the HIB group **(Figure 1C and 1D)**.

As shown in **Figure 2A**, results showed that both the MHCI and MHC II fiber isoforms were expressed in the diaphragm and gastrocnemius muscles. The percentage of type I fibers in the diaphragm and gastrocnemius muscles was significantly increased (33% and 36%, respectively, *P* < 0.05) in the HIB group compared with that in the SA group **(Figure 2B and 2C)**. The proportion of MHC IIa fiber in the HIB group was significantly reduced by 6% (*P* < 0.05) in the diaphragm muscle, while it was significantly increased in the gastrocnemius muscle by 9% (*P* < 0.05) as compared with SA group **(Figure 2B and 2C)**. In contrast, the proportion of MHC IIb fiber was significantly increased by 21% (*P* < 0.05) in the diaphragm muscle, while it was significantly decreased by 15% (*P* < 0.05) in the gastrocnemius muscle of the HIB group as compared with that in the SA group **(Figure 2B and 2C)**. As compared with SA group, the proportion of MHC IIx fiber in the diaphragm and gastrocnemius muscles in the HIB group was significantly increased by 7% (*P* < 0.05) and 16% (*P* < 0.05), respectively **(Figure 2B and 2C)**.

**Figure 2.**
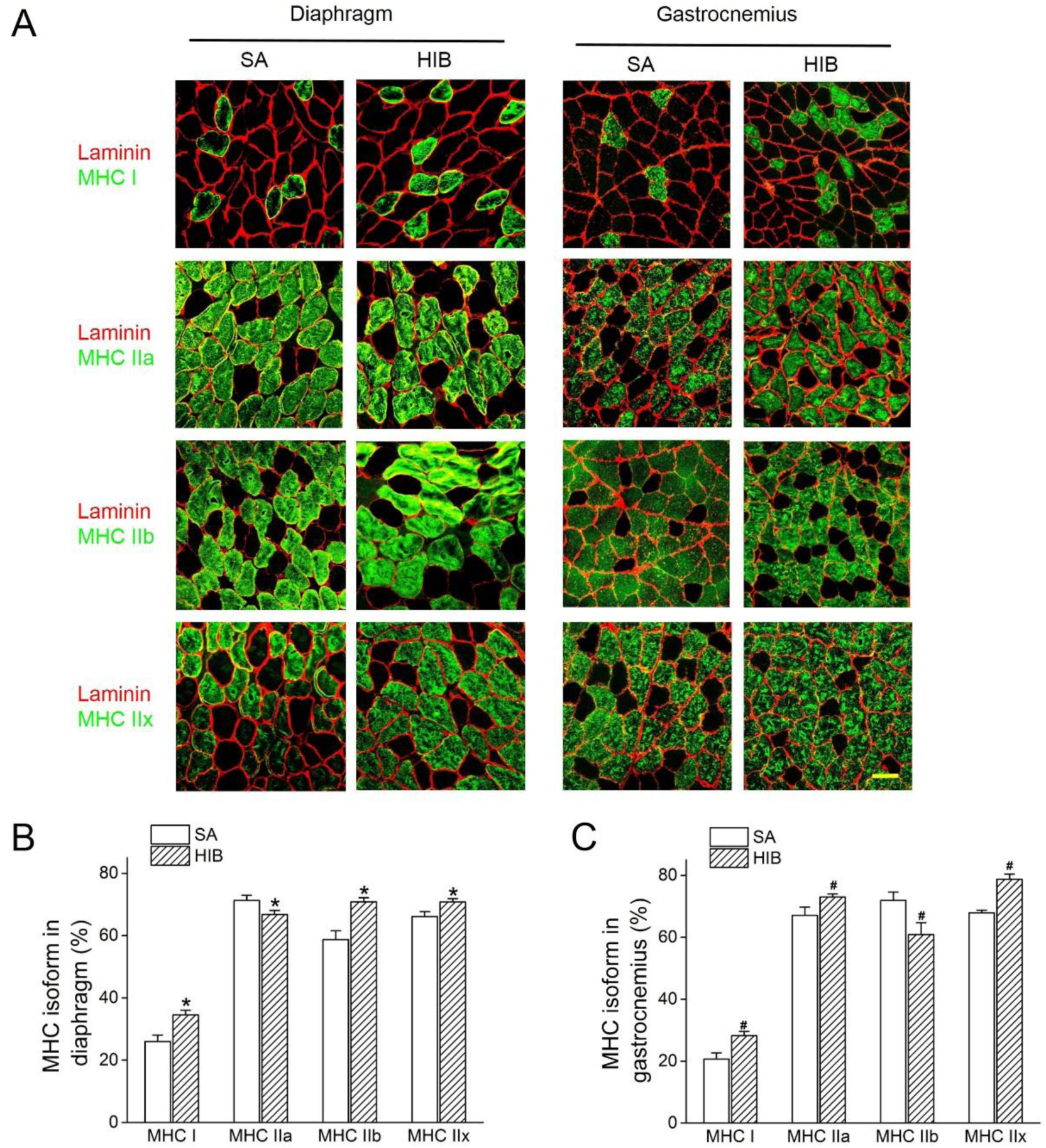
Fiber-type distribution in diaphragm and gastrocnemius muscles. **(A)** Representative immunofluorescent images showing MHC I, MHC IIa, MHC IIb, and MHC IIx fibers with green staining in diaphragm and gastrocnemius muscles. Red represents laminin stain of myofiber interstitial tissue. Scale bar = 50 μm. **(B)** Bar graph showing the changes of fiber-type distribution in diaphragm muscle. **(C)** Bar graph showing the changes of fiber-type distribution in gastrocnemius muscle. SA: summer active ground squirrels, HIB: hibernating ground squirrels. Data represents mean ± SEM, *n* = 8. \**P* < 0.05, compared with SA in diaphragm muscle. ^#^*P* < 0.05, compared with SA in gastrocnemius muscle.

Besides, the protein expression levels of MHC I, MHC IIa, MHC IIb and MHC IIx in the diaphragm and gastrocnemius muscle were analyzed by western blots. As compared with SA group, MHC I protein expression level in the HIB group was significantly increased by 1.41-fold (*P* < 0.05) and 2.07-fold (*P* < 0.05) in the diaphragm and gastrocnemius muscles, respectively **(Figure 3A and 3B)**. In the diaphragm muscle, the protein expression level of MHC IIa was decreased significantly in the HIB group compared with that in the SA group (28%, *P* < 0.05). However, the MHC IIa expression was significantly increased by 1.34-fold (*P* < 0.05) in the gastrocnemius muscle of the HIB group as compared with SA group **(Figure 3A and 3C)**. The protein expression level of MHC IIb was significantly increased by 1.36-fold (*P* < 0.05) in the diaphragm muscle, while it was significantly decreased by 34% (*P* < 0.05) in the gastrocnemius muscle in the HIB group compared with that in the SA group **(Figure 3A and 3D)**. The protein expression level of MHC IIx in the diaphragm and gastrocnemius muscles in the HIB group was 1.48-fold (*P* < 0.05) and 1.3-fold (*P* < 0.05) higher than that in the SA group, respectively **(Figure 3A and 3E)**.

**Figure 3.**
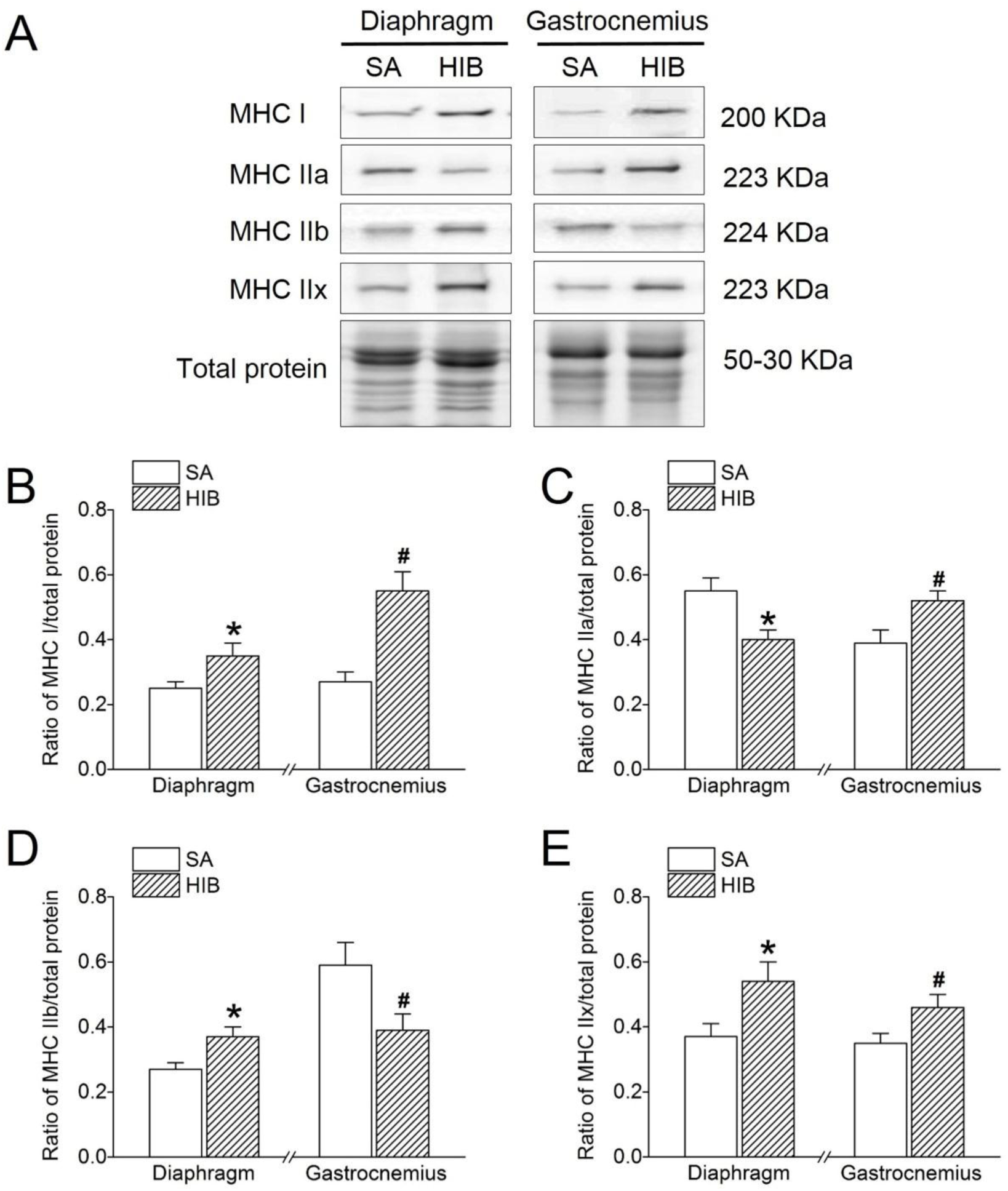
Protein expression levels of MHC I, MHC IIa, MHC IIb, and MHC IIx. **(A)** Representative immunoblots of MHC I, MHC IIa, MHC IIb, and MHC IIx in each group. **(B)** Changes of the ratio of MHC I to total protein. **(C)** Changes of the ratio of MHC IIa to total protein. **(D)** Changes of the ratio of MHC IIb to total protein. **(E)** Changes of the ratio of MHC IIx to total protein. SA: summer active ground squirrels, HIB: hibernating ground squirrels. Data represent mean ± SEM, *n* = 8. \**P* < 0.05, compared with SA in diaphragm muscle. ^#^*P* < 0.05, compared with SA in gastrocnemius muscle.

### Protein expression levels of MuRF1, atrogin-1, Beclin1 and LC3-II indicative of protein degradation metabolism

Compared with that in the SA group, MuRF1 protein expression significantly decreased in the diaphragm muscle (16.0%, *P* < 0.05), but significantly increased in the gastrocnemius muscle (1.3-fold, *P* < 0.05) in the HIB group **(Figure 4A and 4B)**. The expression level of atrogin-1 was significantly decreased by 17% (*P* < 0.05) in the diaphragm muscle, while it was significantly increased by 1.3-fold (*P* < 0.05) in the gastrocnemius muscle in the HIB group compared with that in the SA group **(Figure 4A and 4C)**. In addition, autophagy-related proteins Beclin1 and LC3-II both showed significant increases (1.2-fold and 1.5-fold, respectively, *P* < 0.05) in the diaphragm muscle of the HIB group compared with that in the SA group. In contrast, both were significantly decreased in the gastrocnemius muscle (32.6% and 31.9%, respectively, *P* < 0.05) of the HIB group compared with that of the SA group **(Figure 4A, 4D and 4E)**.

**Figure 4.**
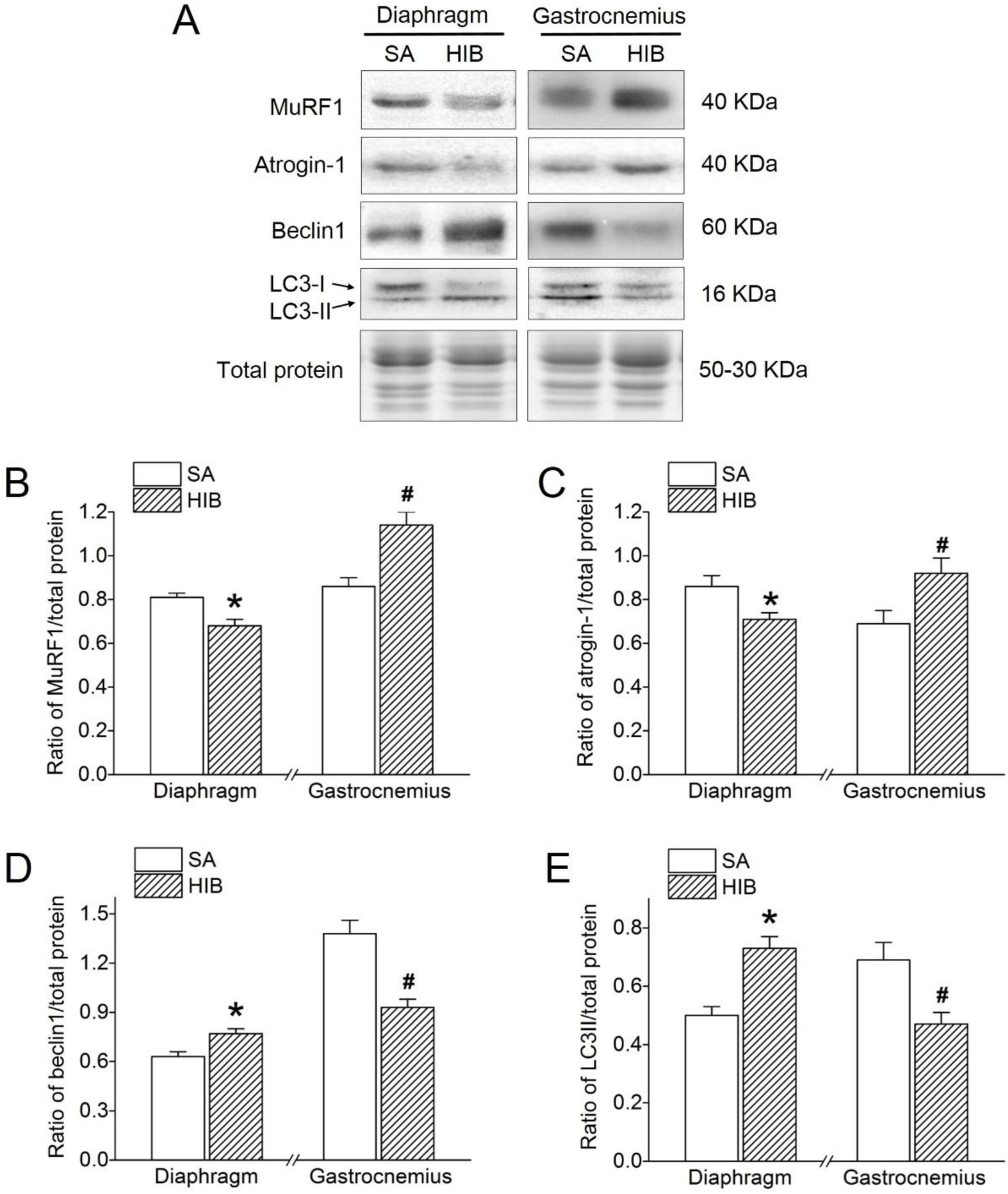
Protein expression levels of MuRF1, atrogin-1, Beclin1 and LC3-II. **(A)** Representative immunoblots of MuRF1, atrogin-1, Beclin1 and LC3-II in each group. **(B)** Changes of the ratio of MuRF1 to total protein. **(C)** Changes of the ratio of atrogin-1 to total protein. **(D)** Changes of the ratio of Beclin1 to total protein. **(E)** Changes of the ratio of LC3-II to total protein. SA: summer active ground squirrels, HIB: hibernating ground squirrels. Data represent mean ± SEM, *n* = 8. \**P* < 0.05, compared with SA in diaphragm muscle. ^#^*P* < 0.05, compared with SA in gastrocnemius muscle.

### Proteasome activity

Compared with SA group, chymotrypsin and trypsin activities in HIB group were significantly decreased by 60.5% and 52.8% (*P* < 0.05), respectively, in diaphragm muscle, but were significantly increased by 32.7% and 79.9% (*P* < 0.05), respectively, in gastrocnemius muscle **(Figure 5A and 5B)**.

**Figure 5.**
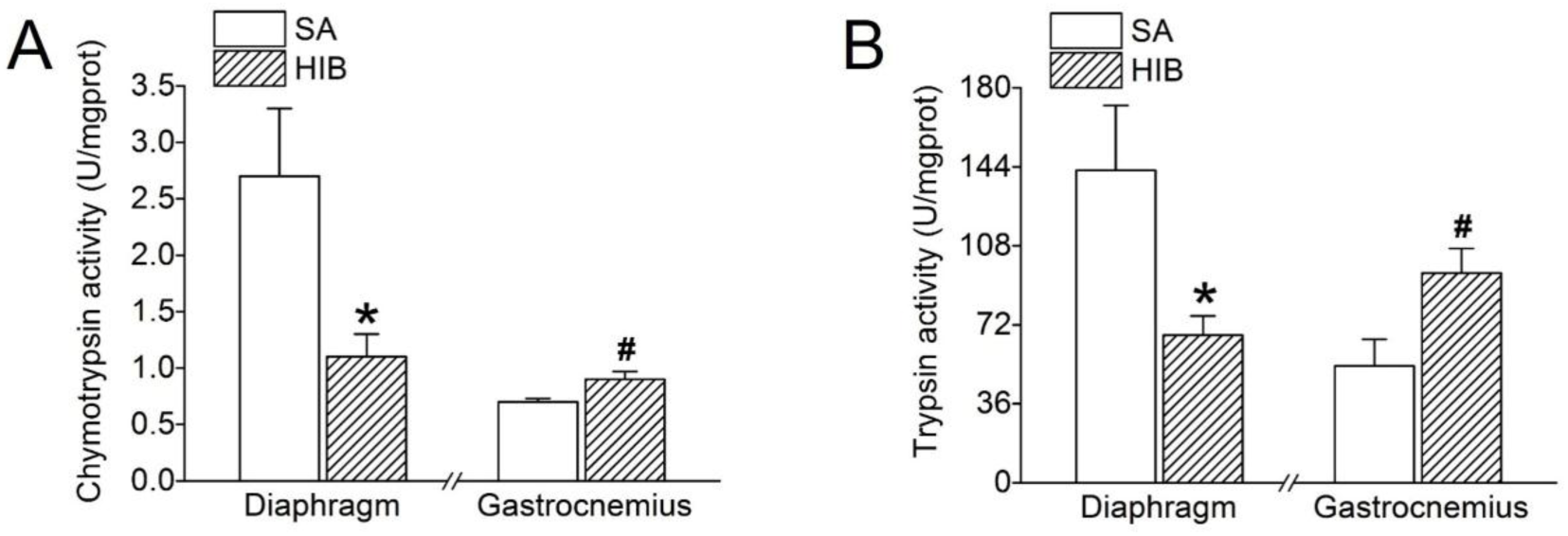
Proteasome activity. **(A) Chymotrypsin proteasome activity. (B) Trypsin proteasome activity**. SA: summer active ground squirrels, HIB: hibernating ground squirrels. Data represent mean ± SEM, *n* = 8. \**P* < 0.05, compared with SA in diaphragm muscle. ^#^*P* < 0.05, compared with SA in gastrocnemius muscle.

### Protein expression levels of calpain-1, calpain-2, calpastatin, desmin and troponin T indicative of protein degradation metabolism

The protein expression level of calpain-1 and calpain-2 both showed a significant decrease (30% and 46.3%, respectively, *P* < 0.05) in the diaphragm muscle of the HIB group compared with levels in the SA group. In contrast, calpain-1 and calpain-2 protein expression levels were significantly increased in the gastrocnemius muscle (1.9-fold and 1.5-fold, respectively, *P* < 0.05) in the HIB group compared with levels in the SA group **(Figure 6A, 6B and 6C)**. Furthermore, contrary to calpain-1 and calpain-2, calpastatin protein expression showed a significant 1.6-fold (*P* < 0.05) increase in the diaphragm muscle, but a significant 32.6% (*P* < 0.05) decrease in the gastrocnemius muscle of the HIB group compared with that in the SA group **(Figure 6A and 6D)**. The protein expression level of desmin was significantly increased by 1.51-fold (*P* < 0.05) in the diaphragm muscle, while it was significantly decreased by 23% (*P* < 0.05) in the gastrocnemius muscle in the HIB group compared with that in the SA group **(Figure 6A and 6E)**. However, the protein expression level of troponin T in both diaphragm and gastrocnemius muscles showed no significant difference between the HIB and SA groups **(Figure 6A and 6F)**.

**Figure 6.**
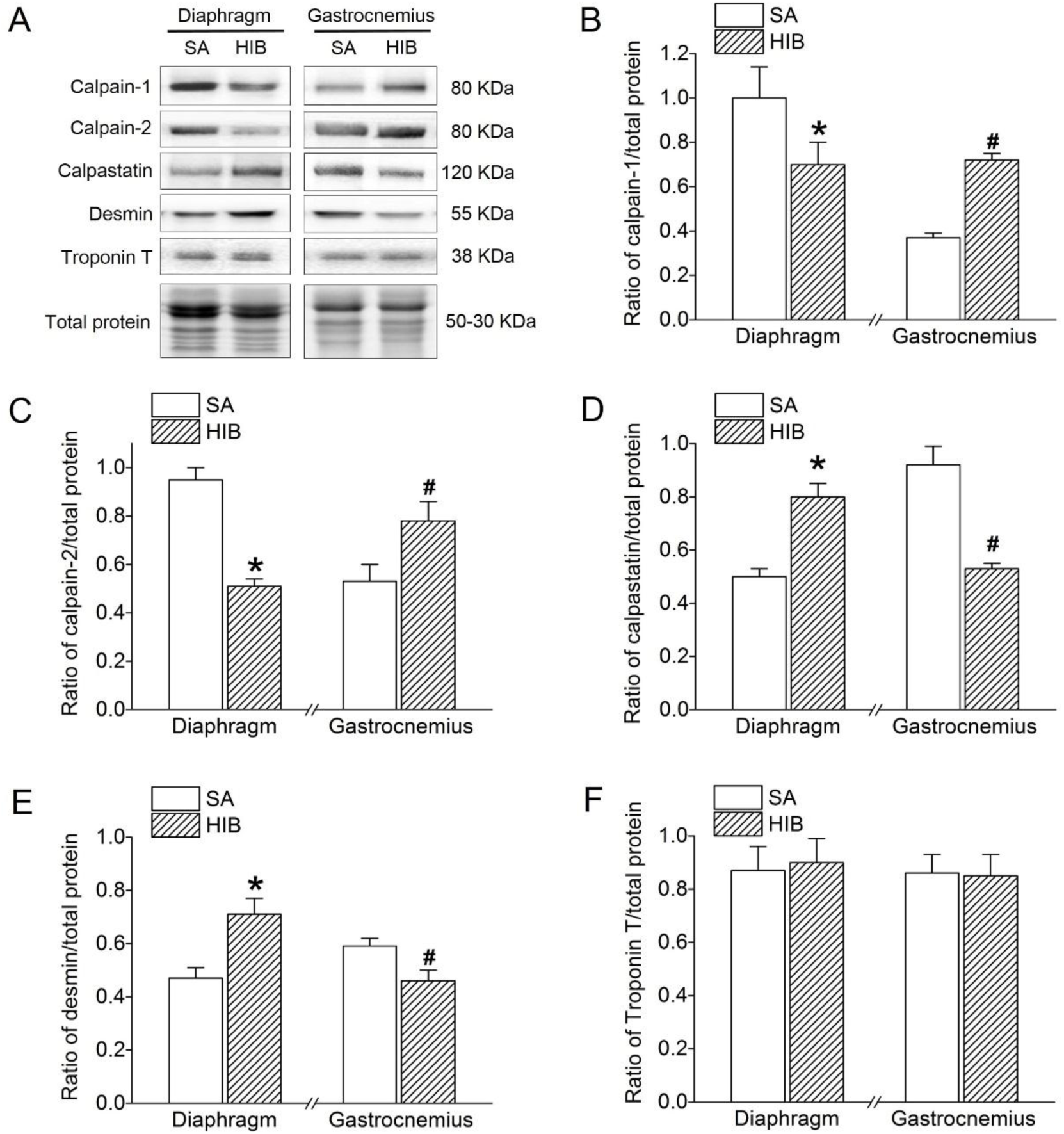
Protein expression levels of calpain-1, calpain-2, calpastatin, desmin and troponin T. **(A)** Representative immunoblots of calpain-1, calpain-2, calpastatin, desmin and troponin T in each group. **(B)** Changes of the ratio of calpain-1 to total protein. **(C)** Changes of the ratio of calpain-2 to total protein. **(D)** Changes of the ratio of calpastatin to total protein. **(E)** Changes of the ratio of desmin to total protein. **(F)** Changes of the ratio of troponin T to total protein. SA: summer active ground squirrels, HIB: hibernating ground squirrels. Data represent mean ± SEM, *n* = 8. \**P* < 0.05, compared with SA in diaphragm muscle. ^#^*P* < 0.05, compared with SA in gastrocnemius muscle.

### Protein expression levels of P-Akt, P-mTORC1, P-S6K1 and P-4E-BP1 indicative of protein synthesis metabolism

Western blots analysis was used to detect the protein expression levels of P-Akt, P-mTORC1, P-S6K1, and P-4E-BP1 in the diaphragm and gastrocnemius muscles. As shown in **Figure 7**, P-Akt protein expression significantly increased by 1.2-fold (*P* < 0.05) in the diaphragm muscle, but significantly decreased by 55.3% (*P* < 0.05) in the gastrocnemius muscle in the HIB group versus the SA group **(Figure 7A and 7B)**. Similarly, P-mTORC1 protein expression showed a significant 1.4-fold (*P* < 0.05) increase in the diaphragm muscle, but a significant 30.9% (*P* < 0.05) decrease in the gastrocnemius muscle in the HIB group compared with that in the SA group **(Figure 7A and 7C)**. Moreover, the P-S6K1 and P-4E-BP1 protein expression levels in the diaphragm muscle increased significantly by 1.4-fold (*P* < 0.05) and 1.3-fold (*P* < 0.05), respectively, in the HIB group compared with that in the SA group. In contrast, the P-S6K1 and P-4E-BP1 protein expression levels in the gastrocnemius muscle decreased significantly by 30.9% (*P* < 0.05) and 22.7% (*P* < 0.05), respectively, in the HIB group compared with levels in the SA group **(Figure 7A, 7D and 7E)**.

**Figure 7.**
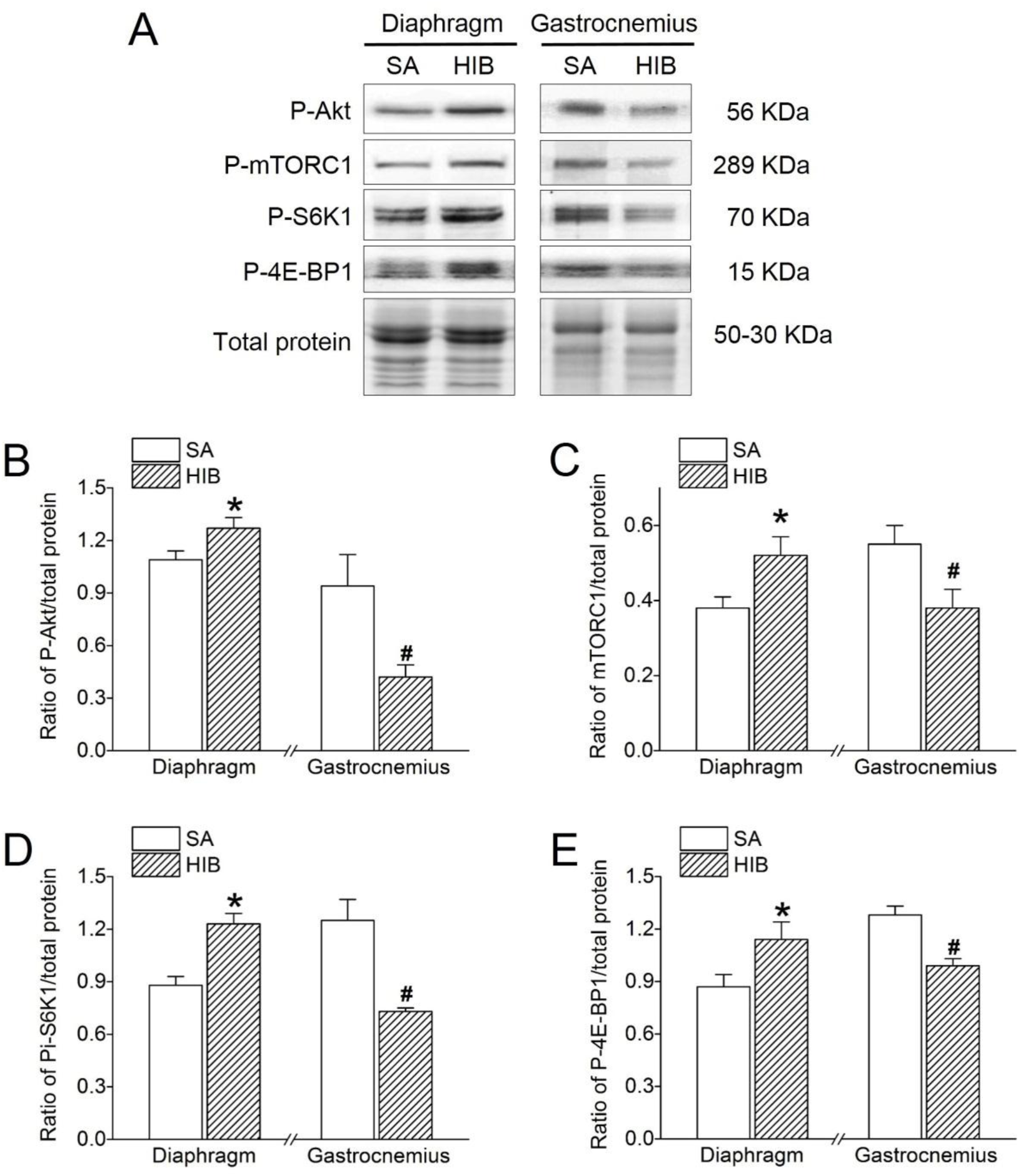
Protein expression levels of P-Akt, P-mTORC1, P-S6K1 and P-4E-BP1. **(A)** Representative immunoblots of P-Akt, P-mTORC1, P-S6K1 and P-4E-BP1 in each group. **(B)** Changes of the ratio of P-Akt to total protein. **(C)** Changes of the ratio of P-mTORC1 to total protein. **(D)** Changes of the ratio of P-S6K1 to total protein. **(E)** Changes of the ratio of P-4E-BP1 to total protein. SA: summer active ground squirrels, HIB: hibernating ground squirrels. Data represent mean ± SEM, *n* = 8. \**P* < 0.05, compared with SA in diaphragm muscle. ^#^*P* < 0.05, compared with SA in gastrocnemius muscle.

### Protein expression levels of MyoD, myogenin and myostatin indicative of muscle regeneration

MyoD and myogenin protein expression levels showed a significant 1.3-fold (*P* < 0.05) and 1.4-fold (*P* < 0.05) increase, respectively, in the diaphragm muscle, but a significant 26.7% (*P* < 0.05) and 12.2% (*P* < 0.05) decrease, respectively, in the gastrocnemius muscle of the HIB group compared with that of the SA group **(Figure 8A, 8B and 8C)**. In contrast, the myostatin protein expression level decreased by 38.2% (*P* < 0.05) in the diaphragm muscle but increased by 2.3-fold (*P* < 0.05) in the gastrocnemius muscle in the HIB group compared with the SA group **(Figure 8A and 8D)**.

**Figure 8.**
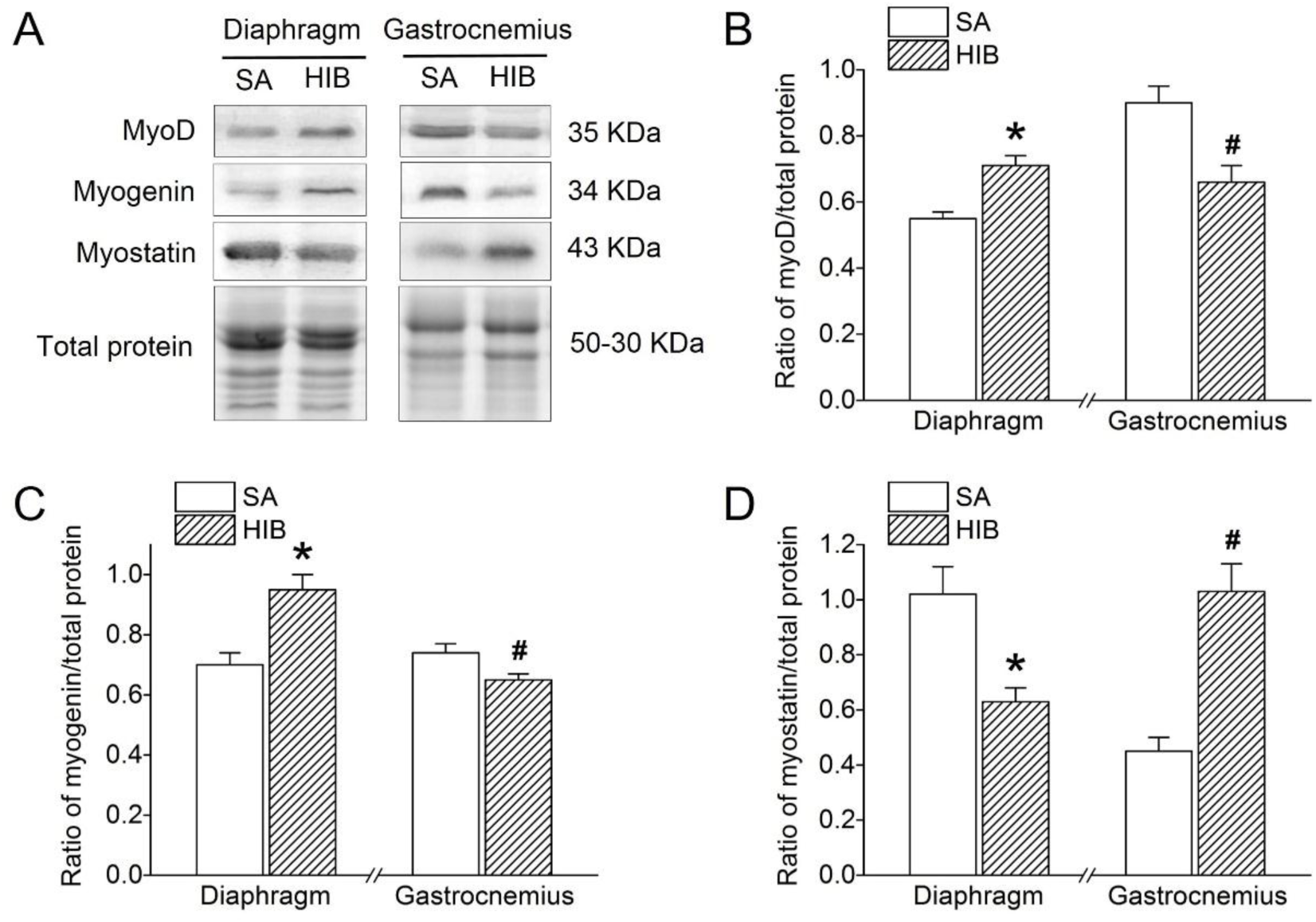
Protein expression levels of MyoD, myogenin and myostatin. **(A)** Representative immunoblots of MyoD, myogenin and myostatin in each group. **(B)** Changes of the ratio of MyoD to total protein. **(C)** Changes of the ratio of myogenin to total protein. **(D)** Changes of the ratio of myostatin to total protein. SA: summer active ground squirrels, HIB: hibernating ground squirrels. Data represent mean ± SEM, *n* = 8. \**P* < 0.05, compared with SA in diaphragm muscle. ^#^*P* < 0.05, compared with SA in gastrocnemius muscle.

## Discussion

In the present study, changes in key signals of protein synthesis, protein degradation, and muscle regeneration were detected to determine the mechanism(s) related to the differential adaptability of the diaphragm and gastrocnemius muscles in hibernating Daurian ground squirrels. The wet weight of the diaphragm muscle was significantly increased, while it was significantly decreased in gastrocnemius muscle **(Figure 1B)**. Due to its importance as a respiratory muscle, the CSA of the diaphragm muscle fibers showed significant hypertrophy (26.1% increase) **(Figure 1C and 1D)** during hibernation. However, the CSA of the gastrocnemius muscle was not completely preserved, instead showing significant atrophy (20.4% decrease) **(Figure 1C and 1D)**. Consistent with our findings, previous studies have also shown significant muscle wet weight loss (14%) in the gastrocnemius muscle of hibernating golden-mantled ground squirrels (Wickler et al. 1991), but significant muscle weight gain (40% and 19%) in the diaphragm muscle of hibernating Syrian hamsters (Deveci and Egginton 2002) and golden-mantled ground squirrels, respectively (Reid et al. 1995). In addition, the results of both immunofluorescent analysis and western blots showed that the expressions of MHC I and MHC IIx in the HIB group were significantly increased in the gastrocnemius and diaphragm muscles, the expression of MHC IIa was significantly increased in the gastrocnemius muscle, while it was significantly decreased in the diaphragm muscle, besides, MHC IIb expression was significantly reduced in the gastrocnemius muscle while it showed a significant increase in the diaphragm muscle compared with the SA group. Furthermore, both the diaphragm and gastrocnemius muscles displayed a significant fast-to-slow fiber type transition **(Figure 2)**. Consistent with our recent study, the gastrocnemius muscle showed a fast-to-slow fiber type transition during hibernation in Daurian ground squirrels (Ma et al. 2019), in contrast, the fiber type in the gastrocnemius muscle of the hindlimb unloading rat showed a significant slow-to-fast fiber type shift (Jones et al. 2003). Both type I and type II fibers of the diaphragm were significantly atrophied in the mechanically ventilated rats (Ichinoseki-Sekine et al. 2014). Therefore, our study again demonstrates that there is a fast-to-slow fiber type transition in different skeletal muscles in hibernating ground squirrels. In this study, we investigated the mechanism(s) related to the differences in the physiological adaptations of the diaphragm and gastrocnemius muscles in hibernating Daurian ground squirrels from three aspects: i.e., protein synthesis, protein degradation, and muscle regeneration.

MuRF1 and atrogin-1 are considered as molecular markers of muscular atrophy. Earlier studies have shown that MuRF1 protein expression is significantly increased in the gastrocnemius muscle of hindlimb-unloaded rats, which subsequently induces overexpression of ubiquitin protein and muscle atrophy (Su et al. 2014). Moreover, the expression level of atrogin-1 mRNA is significantly increased by 2-fold in the gastrocnemius muscle of hindlimb unloading mice (Al-Nassan et al. 2012). Furthermore, both the protein expression levels of MuRF1 and atrogin-1 were significantly increased in the diaphragm of mechanically ventilated rats (Maes et al. 2014). Previous studies have shown that chymotrypsin and trypsin activities were transiently increased in the gastrocnemius muscle after 5 days of hindlimb-unloaded rats (Magne et al. 2011). Earlier studies have reported significant increases in both chymotrypsin and trypsin activity in the diaphragm muscle of mechanically ventilated rats (McClung et al. 2008). Our results showed that MuRF1, atrogin-1 protein expression, chymotrypsin and trypsin activities were all significantly increased in the gastrocnemius muscle of hibernating squirrels compared with the summer active group **(Figure 4B and 4C)**, which is consistent with our recent study (Ma et al. 2019, Wei et al. 2018) and further indicating that high MuRF1 and atrogin-1 protein expression may be involved in protein degradation and gastrocnemius muscle atrophy. In contrast, we found that MuRF1, atrogin-1 protein expression, chymotrypsin and trypsin activities were significantly decreased in the diaphragm muscle, which may be an anti-atrophy mechanism of this muscle in hibernating Daurian ground squirrels.

The calpain system plays an essential role in muscle protein degradation and is a likely component of the targets of disuse atrophy, especially in relation to myofibrillar disassembly (Jackman and Kandarian 2004). Previous studies have demonstrated that calpain-induced myofibrillar protein degradation (Smuder et al. 2010) as well as calpain activity (e.g., by 95%) (Bruells et al. 2016) are both increased in the diaphragm of rats under mechanical ventilation. Our results showed that protein expression of calpain-1 and calpain-2 increased by 94.6% and 47.2%, respectively, whereas that of calpastatin (an endogenous inhibitor of calpains) decreased by 42.4% in the gastrocnemius muscle of the hibernating group compared with that of the summer active group **(Figure 6B, 6C and 6D)**, suggesting that the up-regulation of calpain signals may contribute to gastrocnemius muscle atrophy. However, consistent with our previous studies in the soleus and extensor digitorum longus muscles (both showing no significant atrophy in hibernation) (Chang et al. 2018b, Yang et al. 2014), the protein expression levels of calpain-1 and calpain-2 decreased by 30% and 46.6%, respectively, whereas that of calpastatin increased by 60% in the diaphragm muscle of hibernating ground squirrels compared with the summer active group **(Figure 6B, 6C and 6D)**. Higher calpain-1 (1.44-fold) and calpain-2 (2.38-fold) activity has also been observed in the longissimus dorsi and soleus muscles of hibernating long-tailed ground squirrels (*Spermophilus undulates*) compared with summer active squirrels (Popova et al. 2017). Calpain can hydrolyze substrates such as desmin and troponin T (Barta et al. 2005). Our study showed that desmin was significantly decreased in the gastrocnemius muscle of the HIB group and significantly increased in the diaphragm muscle compared with the SA group, however, troponin T showed no significant changes between the gastrocnemius and diaphragm muscles **(Figure 6)**, which indicated that different substrates have different sensitivity to the calpain system. Our previous study showed desmin expression was unchanged in soleus muscle and extensor digitorum longus muscle of Daurian ground squirrels during hibernation, but was significantly reduced by 39-51% in hindlimb unloading rats (Chang et al. 2018b). Our recent study also showed that the protein expression of desmin in gastrocnemius muscle was significantly decreased, while troponin T was unchanged during hibernation in Daurian ground squirrels (Ma et al. 2019). Therefore, our data suggest that the muscle-specific activation of the calpain system and degradation of different substrates may be involved in the different adaptabilities of the diaphragm and gastrocnemius muscles in Daurian ground squirrels during hibernation.

Beclin1 (B-cell Lymphoma 2 interacting protein 1) and LC3 (microtubule-associated protein light chain 3) are two molecular markers of autophagy. Beclin1 can form complexes that induce proteins to be located on the autophagic membrane (Kihara et al. 2001). Furthermore, LC3 is involved in the formation of the autophagic membrane, whereby LC3-I converts into LC3-II for the formation of autophagosomes (Tanida et al. 2005, Tanida et al. 2004). Earlier studies have reported significant increases in the mRNA and protein expression levels of Beclin1 (1.91-fold and 30%, respectively) and LC3 (1.82-fold and 30%, respectively) in the diaphragm muscles of mechanically ventilated rats (Smuder et al. 2018). Furthermore, hindlimb unloading in mice reportedly results in the up-regulation of Beclin1 mRNA and induction of autophagy in soleus muscle atrophy (Cannavino et al. 2014). In the current study, the protein expression levels of Beclin1 and LC3-II in the gastrocnemius muscle of hibernating ground squirrels were significantly lower (Beclin1, −31.9%, LC3-II, −32.6%, *P* < 0.05) than those in the summer active group, whereas the expression levels of Beclin1 and LC3-II in the diaphragm muscle were significantly higher (Beclin1, 46%, LC3-II, 22.2%, *P* < 0.05). Our results showed that autophagy was suppressed in the gastrocnemius muscle but was promoted in the diaphragm muscle of the hibernating ground squirrels. Consistent with our gastrocnemius muscle results, earlier research demonstrated a decrease in the LC3-II/LC3-I ratio in the quadriceps and tibialis anterior muscles of thirteen-lined ground squirrels during hibernation, thus indicating suppression of autophagy (Andres-Mateos et al. 2013). Our data suggest that protein degradation causing gastrocnemius muscle atrophy occurred primarily through the ubiquitin pathway rather than the autophagy-lysosome pathway. The elevated level of autophagy in the diaphragm muscle may correspond to the elevated level of diaphragm muscle protein synthesis, both of which promoted protein turnover and diaphragm hypertrophy.

As key molecules mediating the Akt-mTOR signaling pathway in protein synthesis, the protein expression levels of Akt and mTOR are significantly decreased in the diaphragm after mechanical ventilation, as is the protein expression of downstream target 4E-BP1 (Hudson et al. 2015, Yu et al. 2018). Moreover, gastrocnemius myofibrillar protein content and synthesis (mg/day) are shown to decrease significantly (by 26% and 64%, respectively) in rats undergoing tail suspension (Linderman et al. 1994). In the present study, four protein synthesis markers, i.e., P-Akt, P-mTORC1, P-S6K1, and P-4E-BP1, were significantly increased by 17%, 37%, 40%, and 31% (*P* < 0.05), respectively, in the diaphragm muscle, but were significantly decreased by 55%, 31%, 42% and 23% (*P* < 0.05), respectively, in the gastrocnemius muscle during hibernation compared with levels during summer activity **(Figure 7)**. Similar to our findings in the gastrocnemius muscle, Akt activity and phosphorylated Akt (Ser 473) are reported to decrease (by 60% and 40%, respectively) in skeletal muscle during hibernation in Richardson’s ground squirrels (*Spermophilus richardsonii*) compared with euthermic controls, which may contribute to the coordinated suppression of energy-expensive anabolism during winter torpor (Abnous et al. 2008). Moreover, research has shown that Akt/PKB activity exhibits seasonal peaks in the gastrocnemius muscle of yellow-bellied marmots (*Marmota flaviventris*) during their annual cycle, thereby demonstrating tissue-specific seasonal activation (Hoehn et al. 2004). Thus, our results suggest that the up-regulation of protein synthesis contributes to the diaphragm muscle hypertrophy, whereas the down-regulation of protein synthesis contributes to the gastrocnemius muscle atrophy found in hibernating Daurian ground squirrels.

Whether muscle regeneration potential in different skeletal muscles is altered in response to hibernation has remained unexplored. Here, we determined the protein expression levels of key markers, such as MyoD, myogenin, and myostatin, in satellite cell regulatory pathways. The expression level of MyoD showed a significant 1.3-fold (*P* < 0.05) increase in the diaphragm muscle, but a significant 26.7% (*P* < 0.05) decrease in the gastrocnemius muscle of hibernating squirrels compared with the active group **(Figure 8A and 8B)**. Mechanical ventilation is known to induce a decrease in MyoD mRNA and protein expression in the diaphragm muscle of rats (Maes et al. 2008, Racz et al. 2003), suggesting the inactivation of quiescent satellite cells. Furthermore, muscle disuse can induce a decline in satellite cells together with an up-regulation in MyoD, suggesting that activation of satellite cells and fusion to existing myofibers may take place during hindlimb immobilization (Guitart et al. 2018). Unlike our results, however, earlier research found no changes in MyoD protein levels, but a decrease in mRNA transcript levels in several hindlimb thigh muscles of thirteen-lined ground squirrels during hibernation (Tessier and Storey 2010). Therefore, our data suggest that the differential regulation of MyoD protein expression in muscle regeneration may be involved in the different adaptabilities of the diaphragm and gastrocnemius muscles in ground squirrels.

Previous studies have found that myogenin protein content is severely decreased (e.g., by 63%) in the diaphragm muscle after mechanical ventilation (Maes et al. 2008). Furthermore, although myogenin mRNA expression showed no change in 7-d immobilized mice (Guitart et al. 2018), levels were found to double (*P* < 0.05) following two weeks of immobilization in healthy young men (Wall et al. 2014). In the current study, the expression level of myogenin showed a significant 1.4-fold (*P* < 0.05) increase in the diaphragm muscle, but a significant 12.2% (*P* < 0.05) decrease in the gastrocnemius muscle of the hibernating squirrels compared with the summer active squirrels **(Figure 8A and 8C)**. Consistent with our diaphragm muscle results, myogenin is reported to be up-regulated (2.4-fold) in the cardiomyocytes of thirteen-lined ground squirrels during late torpor, indicating a propensity for hypertrophy (Zhang et al. 2016a). Furthermore, in accordance with our gastrocnemius muscle results, myogenin expression is significantly decreased (by 52%) in the hindlimb thigh muscles, including oxidative and glycolytic fiber types, of thirteen-lined ground squirrels during late torpor relative to the euthermic state (Zhang et al. 2016b). Therefore, we speculate that high myogenin expression contributes to diaphragm hypertrophy, whereas low expression contributes to gastrocnemius atrophy in hibernating Daurian ground squirrels.

Myostatin is a potent negative regulator of skeletal muscle growth. Moreover, myostatin negatively regulates the activity of the Akt pathway, which promotes protein synthesis and increases activity of the ubiquitin-proteasome system to induce atrophy(Rodriguez et al. 2014). Up-regulation of myostatin can be triggered by mechanical ventilation (Liang et al. 2018). Furthermore, myostatin knockout-induced hypertrophy involves little or no input from satellite cells (Amthor et al. 2009). However, the role of myostatin in disuse muscle atrophy remains controversial. For example, increased myostatin mRNA and protein levels have been reported in mice following 11-d of spaceflight (Allen et al. 2009) and in humans after chronic disuse (Reardon et al. 2001); in contrast, decreased myostatin protein levels have been observed after 14 d of immobilization in healthy young men (Wall et al. 2014). In the current study, the protein expression level of myostatin decreased by 38.2% (*P* < 0.05) in the hypertrophic diaphragm muscle but increased by 2.3-fold (*P* < 0.05) in the atrophic gastrocnemius muscle **(Figure 8A and 8D)**. Research thus far has indicated that myostatin protein levels are maintained in the hindlimb thigh muscles of thirteen-lined ground squirrels during early hibernation and torpor (Brooks et al. 2011). Inconsistent with our findings, thirteen-lined ground squirrels show low levels of myostatin in the gastrocnemius muscle over prolonged hibernation, indicating possible slow regeneration in response to injury and possible protection against the formation of fibrotic tissue (Andres-Mateos et al. 2012). Other research has also shown unaltered myostatin expression in atrophied left ventriculars of grizzly bears (*Ursus arctos horribilis*), who tolerate extended periods of extremely low heart rate during hibernation (Barrows et al. 2011). Therefore, our results suggest different regulation mechanisms of muscle regeneration in the diaphragm and gastrocnemius muscles of hibernating animals, whereby the muscle regeneration potential of the diaphragm is enhanced and that of the gastrocnemius muscle is weakened. Variations in muscle regeneration ability may participate in the different adaptive mechanisms of muscles in hibernators. A major novelty in the current study was our use of two different types of skeletal muscle to clarify muscle regeneration potential in hibernation.

In conclusion, different patterns of protein synthesis, protein degradation, and muscle regeneration were demonstrated in gastrocnemius muscle atrophy and diaphragm muscle hypertrophy during hibernation. Specifically, in the gastrocnemius muscle, protein synthesis, autophagy, and muscle regeneration were suppressed, and protein degradation was promoted, which likely contributed to muscle atrophy. In contrast, in the diaphragm muscle, protein synthesis, autophagy, and muscle regeneration were promoted, and protein degradation was inhibited, which likely contributed to muscle hypertrophy. Thus, the changes in protein synthesis and degradation displayed muscle specificity during hibernation. Collectively, the differences in muscle regeneration potential and regulatory patterns of protein metabolism may contribute to the different adaptabilities of the diaphragm and gastrocnemius muscles in Daurian ground squirrels during hibernation.

## Abbreviation

Akt: protein kinase B
mTOR: the mammalian target of rapamycin
mTORC1: mammalian target of rapamycin in complex 1
S6K1: 70 kDa ribosomal protein S6 kinase 1
4E-BP1: eukaryotic initiation factor 4E-binding protein 1
CSA: cross-sectional area
myoD: myogenic differentiation 1
Beclin1: B-cell Lymphoma 2 interacting protein 1
LC3: microtubule-associated protein light chain 3
MuRF1: Muscle RING finger 1
Atrogin-1/MAFbx: muscle atrophy F-box

## Acknowledgements

This study was supported by funds from the National Nature Science Foundation of China (31640072), National Nature Science Foundation of China (No. 31772459).

